# SeqTailor: a user-friendly webserver for the extraction of DNA or protein sequences from next-generation sequencing data

**DOI:** 10.1101/408625

**Authors:** Peng Zhang, Bertrand Boisson, Jean-Laurent Casanova, Laurent Abel, Yuval Itan

## Abstract

Human whole-genome sequencing generally reveals about 4,000,000 genetic variants, including 20,000 coding variants, in each individual studied. These data are mostly stored as VCF-format files. Although many variant analysis methods accept VCF files as input, many other tools require DNA or protein sequences, particularly for splicing prediction, sequence alignment, phylogenetic analysis, and structure prediction. However, there is currently no existing online tool for extracting DNA or protein sequences for genomic variants from VCF files with user-defined parameters in a user-friendly, efficient, and standardized manner. We developed the SeqTailor webserver to bridge this gap. It can be used for the rapid extraction of (1) DNA sequences around genetic variants, with customizable window sizes, from the hg19 or hg38 human reference genomes; and (2) protein sequences encoded by the DNA sequences around genetic variants, with built-in SnpEff annotation and customizable window sizes, from human canonical transcripts. The SeqTailor webserver streamlines the sequence extraction process, and accelerates the analysis of genetic variant data with software requiring DNA or protein sequences. SeqTailor will facilitate the study of human genomic variation, by increasing the feasibility of sequence-based analysis and prediction. The SeqTailor webserver is freely available from http://shiva.rockefeller.edu/SeqTailor/.

## INTRODUCTION

Human whole-genome sequencing (WGS) and whole-exome sequencing (WES) generate large amounts of genetic data, generally revealing about 4,000,000 variants (WGS) and 20,000 coding variants (WES), in each individual studied. These variant data are usually recorded in variant call format (VCF), which includes structured fields for chromosome (CHROM), position in reference genome sequence (POS), reference allele (REF), alternative allele (ALT) and others (e.g. QUAL, FILTER, INFO, etc.). The major human genetic variation and disease databases (e.g. gnomAD/ExAC (Lek et al. 2016), 1,000 Genomes Project (Genomes Project et al. 2015), BRAVO (TOPMed 2016), HGMD (Stenson et al. 2012), and ClinVar (Landrum et al. 2014)), and most research laboratories and publications, present and manage genetic variants in VCF. For further variant annotations, predictions and visualizations, most of the leading tools use VCF as input (e.g. ANNOVAR (Wang et al. 2010), SnpEff (Cingolani et al. 2012), VEP (McLaren et al. 2016), CADD (Kircher et al. 2014), and PopViz (Zhang et al. 2018)).

However, as the DNA sequence around the variant site presents richer information than the variant alone, many popular genomic software suites require DNA sequences as input, particularly for predictions relating to splicing (e.g. Human Splicing Finder (Desmet et al. 2009), NetGene2 (Brunak et al. 1991), Spliceman (Lim and Fairbrother 2012), RegRNA (Chang et al. 2013)), sequence alignment with reference genomes (e.g. BLAST (Altschul et al. 1997), BLAT (Kent 2002)), alignments of multiple sequences supplied by the user (e.g. MUSCLE (Edgar 2004), MAFFT (Katoh et al. 2017)), and phylogenetic analyses (e.g. PAML (Yang 2007), IQ-TREE (Trifinopoulos et al. 2016)). In addition, many protein bioinformatics tools use amino-acid sequences for protein domain identification and functional annotation (e.g. Pfam (Finn et al. 2016), InterPro (Finn et al. 2017), PolyPhen-2 (Adzhubei et al. 2010)), protein structure prediction and homology modeling (SWISS-MODEL (Waterhouse et al. 2018), HMMER (Potter et al. 2018)), and protein feature calculation (e.g. POSSUM (Wang et al. 2017), PROFEAT (Zhang et al. 2017)). The sequence alignment tools cited above can also be applied to protein sequences. Hence, a knowledge of the reference protein sequence and of the protein sequences altered by genomic variants would make it possible to evaluate their potential effects on protein domains, structures, and functions.

Users wishing to analyze VCF data with software that requires DNA or protein sequence must therefore extract the corresponding sequence from VCF files. The major tools currently available for this purpose are BEDTools (Quinlan and Hall 2010), GATK (McKenna et al. 2010), and IGV software (Thorvaldsdottir et al. 2013). However, BEDTools and GATK are script-based methods that require programming expertise, and IGV is impractical for high-throughput applications (a detailed feature comparison is presented in the Discussion section). We therefore developed the SeqTailor webserver, which offers a user-friendly, efficient, and standardized approach for streamlining DNA and protein sequence extraction from the genetic variant data in VCF.

## RESULTS

We developed the SeqTailor webserver to provide a user-friendly and efficient approach for the rapid extraction of DNA and protein sequences for genetic variants (single nucleotide variants (SNVs) and indels) in VCF, with user-defined window sizes. It accepts input from either the webpage textbox or a user-uploaded file, and it outputs the extracted sequences in the browser and in a downloadable FASTA file. SeqTailor bridges the gap between genomic variants and the tools requiring DNA or protein sequences, as illustrated in **Figure 1.** The SeqTailor interface is shown in **Figure 2**, and a schematic diagram is provided in **Figure 3**.

**Figure 1:**
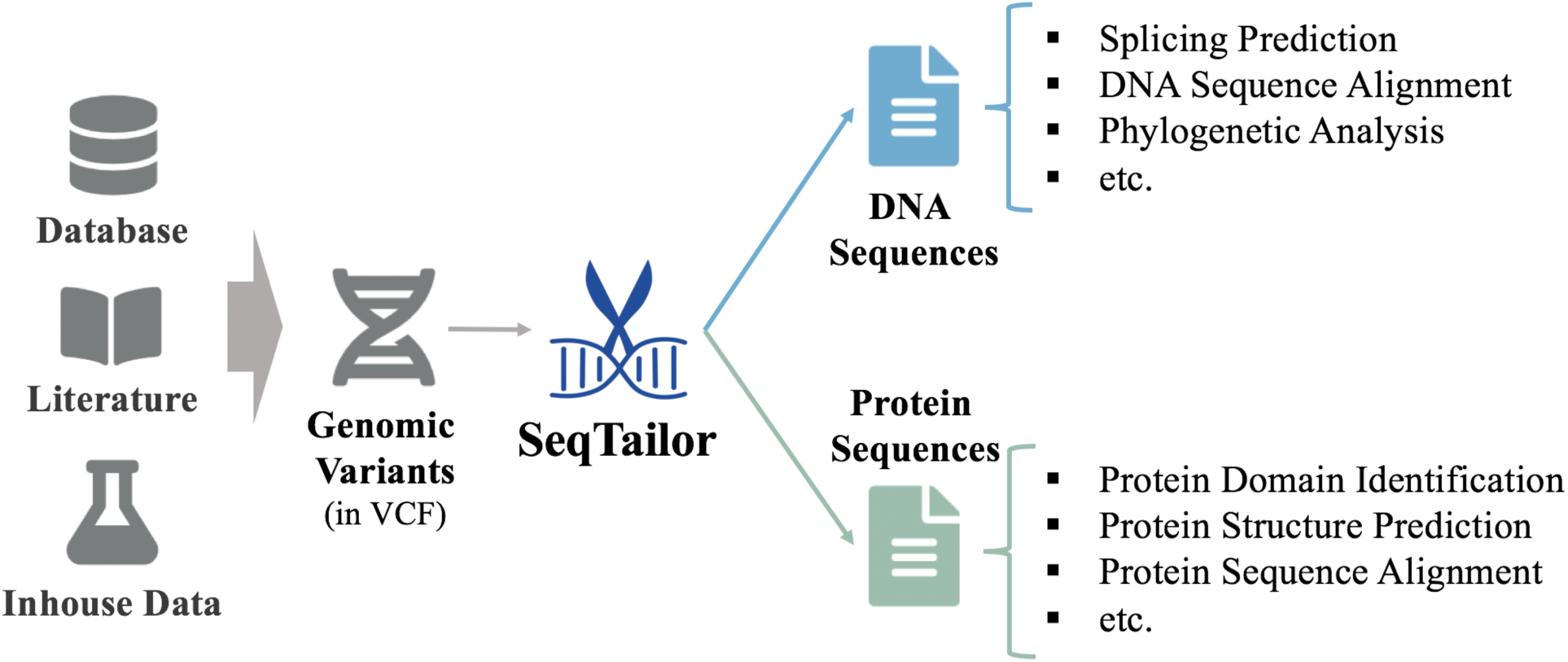
The SeqTailor webserver bridges the gap between genomic variants and sequence-based tools.

**Figure 2:**
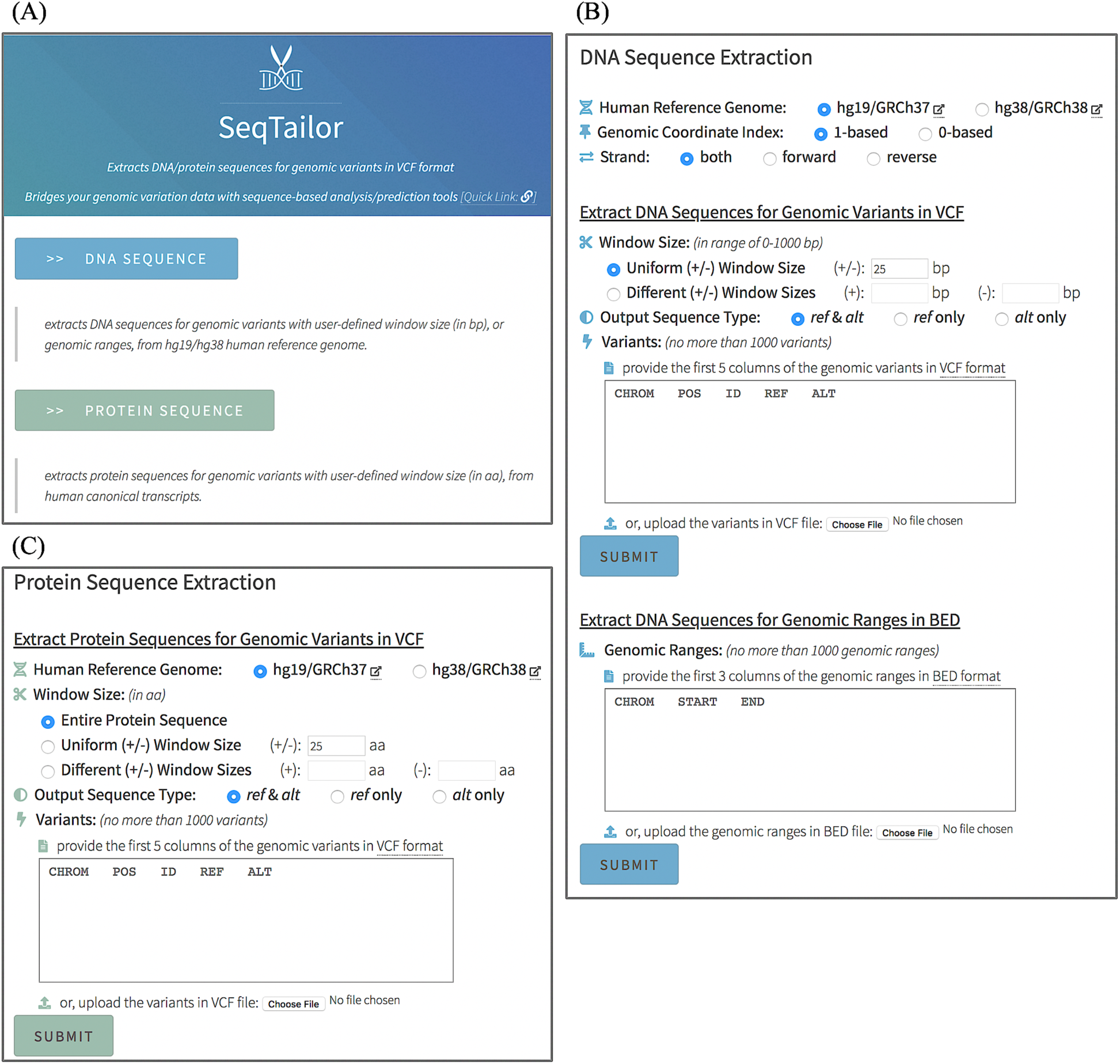
SeqTailor webserver interface: (A) the homepage, (B) the page for DNA sequence extraction, and (C) the page for protein sequence extraction.

**Figure 3:**
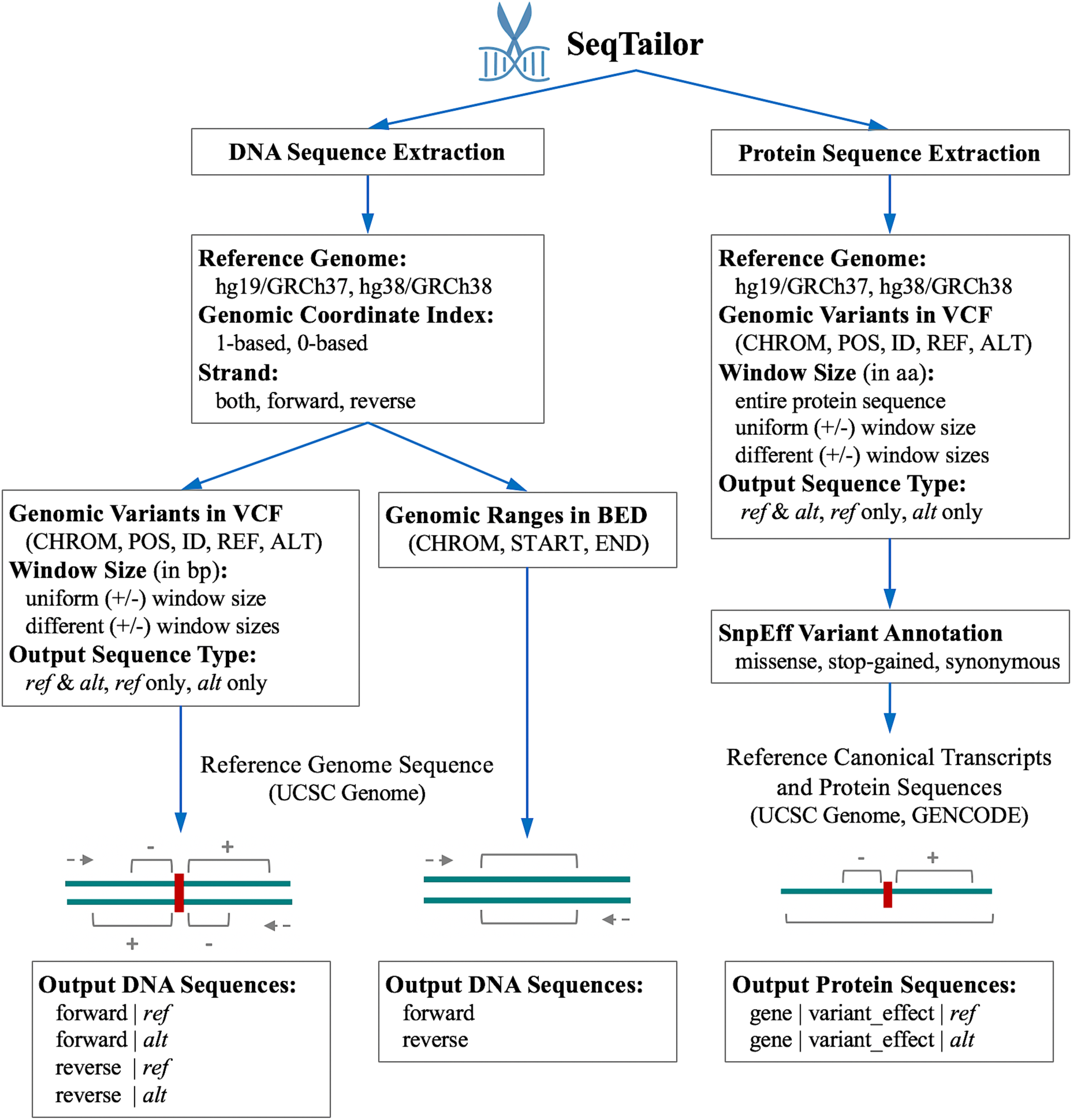
SeqTailor webserver scheme to demonstrate the framework of the application.

For DNA sequence extraction, SeqTailor supports use of the hg19/GRCh37 and hg38/GRCh38 human reference genomes from the UCSC genome database (Karolchik et al. 2004), 1/0-based genomic coordinate indexing, and the choice of the both/forward/reverse strand(s). It accepts both genomic variants in VCF and genomic ranges in BED. For input in the form of genomic variants in VCF, it outputs the wild-type reference (ref) and mutated alternative (alt) DNA sequences, with user-defined up/downstream window sizes (in bp), on the selected strand(s). For input in the form of genomic ranges in BED, it outputs the DNA sequences within the intervals on the selected strand(s). SeqTailor provides the DNA sequences exactly from the START position to the END position, whereas the UCSC Genome Browser always provides an overlap between the input genomic intervals and certain genomic regions (exons, introns, UTRs, etc.), and outputs the desired DNA sequence intervals segregated into different regions.

For protein sequence extraction, SeqTailor first conducts built-in SnpEff variant annotation (Cingolani et al. 2012) for the input variants, retains the missense, stop-gained, and synonymous variants, and extracts the annotated gene symbols, transcript IDs, and amino-acid changes. Only the annotations on canonical transcripts are retained, by matching against the canonical transcripts collected from the UCSC Genome Browser and the GENCODE database (Harrow et al. 2012). SeqTailor provides options for extracting the entire protein sequence, or the sequence within a window of customized size (in aa). Finally, it outputs the canonical and altered protein sequences. As the extraction of protein sequences from genomic ranges is already well-handled by UCSC Genome Browser, we did not develop this function in SeqTailor.

Four case studies are presented in the **Supplementary Document**, including two examples of DNA sequence extraction for a SNV and an indel from the *MSH2* and *BRCA2* genes, respectively, followed by splicing prediction; one example of protein sequence extraction for a stop-gained SNV in the *CFTR* gene; and one example of protein sequence extraction for a SNV in the *BRAF* gene, followed by protein domain search and functional prediction. These examples demonstrate the practical power of SeqTailor to bridge the gap between genome variant data and sequence-based tools for downstream analysis and prediction. SeqTailor will make it efficient to investigate genomic variant data further and will render sequence-based software more accessible.

SeqTailor is designed for the rapid computational extraction of DNA and protein sequences from genetic variant data. For DNA sequence extraction, it takes about 6, 12, 13 and 14 seconds to extract the sequences of 10, 100, 1,000, and 10,000 genomic variants, respectively, from VCF files. For protein sequence extraction, all the input variants are first passed through the build-in SnpEff annotator, and only the missense, stop-gained, and synonymous variants annotated on canonical transcripts are retained for the extraction of protein sequences. It takes approximately 50, 60, 90, 100 seconds to complete the variant annotation/filtering and the protein sequence extraction for 10, 100, 1,000, and 10,000 variants, respectively. The SeqTailor runtime performance is shown in **Figure 4**.

**Figure 4:**
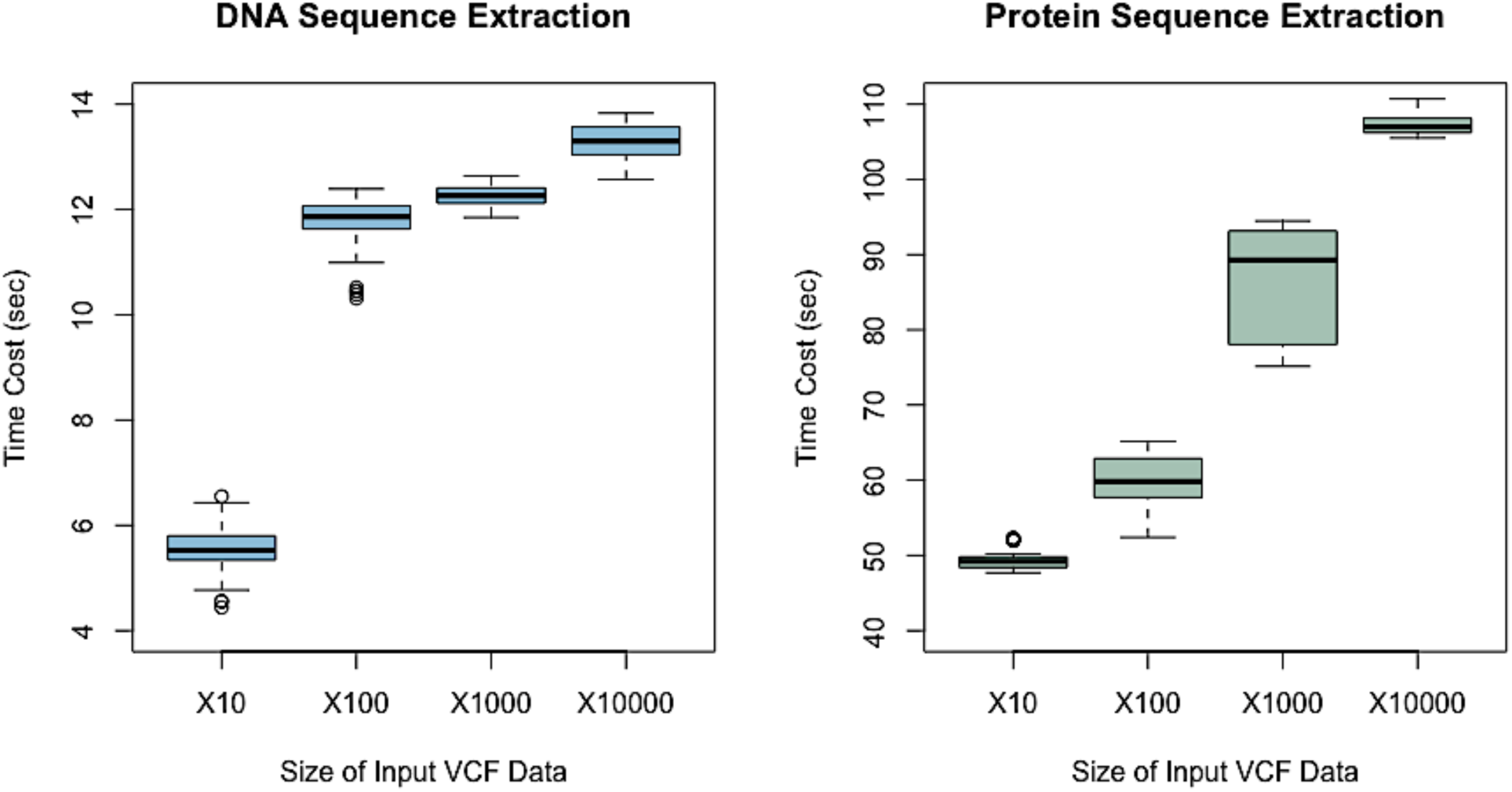
Runtime performance of SeqTailor for extracting DNA or protein sequences from varying sizes of input VCF data (10, 100, 1,000, and 10,000 genomic variants)

## DISCUSSION

Two major sequence extraction methods currently available, BEDTools and GATK, provide a script-based option for DNA sequence extraction, and require significant bioinformatics expertise, and they do not support the extraction of protein sequences around genomic variants. Alternatively, IGV software can be used to locate the position of the variant in the reference genome, with manually copying and pasting to extract the desired DNA sequences or the corresponding protein sequences, one at a time, which would be very time-consuming, error-prone, and impractical in high-throughput data applications. The UCSC Genome Browser provides a web interface for converting genomic ranges in BED/position format to DNA/protein sequences, but users need to go through several steps and define a range of parameters (annotation track, gene table, field table, retrieval region, etc.) to get the result. Importantly, the UCSC Genome Browser does not support sequence extraction from genomic variant data in VCF. The popular GALAXY platform (Afgan et al. 2018) and biopython library (Cock et al. 2009) do not offer options for DNA and protein sequence extraction. A comparison of these approaches with SeqTailor is presented in **Table 1.**

**Table 1:**
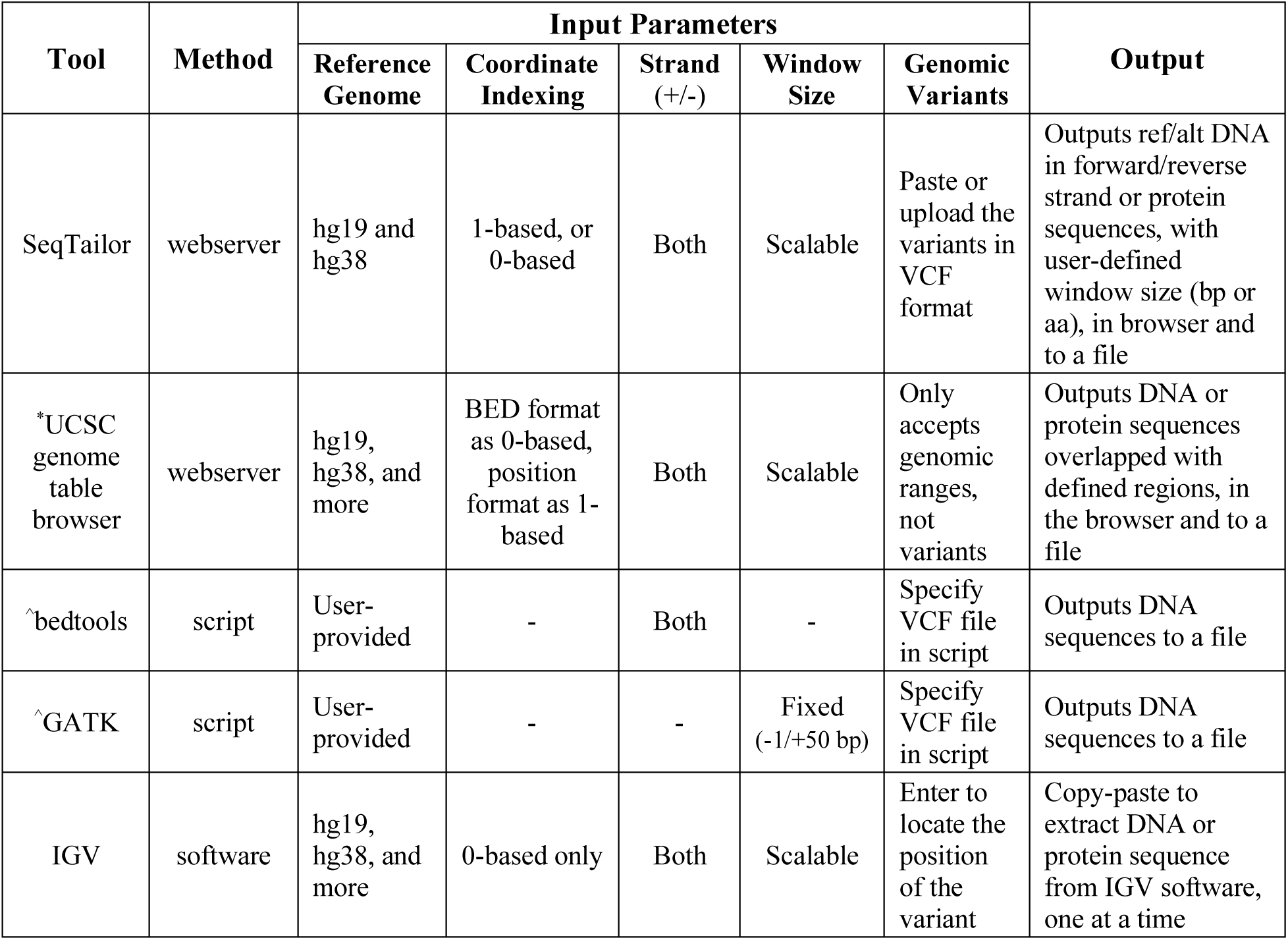
Existing approaches to the extraction of DNA/protein sequences from genomic variants in VCF (* UCSC genome table browser does not accept variants in VCF, but does accept genomic ranges in BED/position format; ^ bedtools and GATK support only DNA sequence extraction, not protein sequence extraction)

SeqTailor webserver is the first online tool for the extraction of DNA and protein sequences from genomic variants in VCF. It presents an efficient and customizable approach for this purpose, requiring only succinct input parameters and no knowledge of script programming. It will be of benefit to scientists studying human genetics, genomics, and molecular biology, enabling them to perform further analyses of their variants with DNA or protein sequence-based tools more rapidly and efficiently than is currently possible.

## METHODS

### Webserver

The SeqTailor webserver is run by Apache HTTP (version 2.2.15), on a Red Hat Enterprise Linux Server (version 6.9), with 8 Intel CPU processors @ 2.4GHz and 48GB RAM. The website interface is design and presented in HTML, PHP, CSS, and JavaScript. The data are stored in MySQL tables and plain text files. The sequence extraction programs are coded in Python.

### Reference Genome

The human reference genome sequences (GRCh37/hg19 assembly, and GRCh38/hg38 assembly) are obtained from UCSC Genome Browser, and stored by genome and by chromosome, to accelerate data processing. Once SeqTailor receives the variants for DNA sequence extraction, it firstly groups the variants by chromosome, and then extracts all the DNA sequences from the same chromosome only once, thus enhancing the efficiency of sequence extraction.

### Canonical Transcript

The canonical transcript IDs were retrieved from UCSC Genome Browser, and the protein-coding transcript translated sequences were downloaded from the GENCODE database. Merging these two resources of data, we obtained a collection of amino-acid sequences for 17,675 human canonical transcripts.

### Built-in Variant Annotation

SnpEff is built in for variant annotation and effect prediction within SeqTailor, with the installed GRCh37.75 and GRCh38.85 databases. Its speed is maximized by a multi-thread configuration, annotating the canonical transcript only, and excluding the upstream, downstream, intergenic, and intron regions. The input variants are then filtered by annotated effect, for retention of the missense, stop-gained, and synonymous variants only. The remaining transcripts are matched against the canonical transcript sequences collected from UCSC genome browser and GENCODE. The annotated amino-acid changes are then reflected in the canonical protein sequences.

### Runtime Evaluation

The runtime evaluation was performed by applying four input data sizes (10, 100, 1,000, and 10,000 genetic variants). We generated 50 VCF files for each input size, by randomly selected variants from the gnomAD database (Lek et al. 2016), and retaining only the first five columns of information (CHROM, POS, ID, REF, and ALT). We therefore used a total of 200 VCF files with different numbers of variants to evaluate the runtime performance of SeqTailor for DNA and protein sequence extraction.

## ACKNOWLEDGEMENT

We thank Y. Nemirovskaya and D. Papandrea for administrative support, F. Rapaport and B. Bigio for technical discussions, and Z. Yang for the artwork of website design. This study was supported by the Rockefeller University, and the Charles Bronfman Institute for Personalized Medicine, Icahn School of Medicine at Mount Sinai.

